# Retrosplenial and hippocampal synchrony during retrieval of old memories in macaques

**DOI:** 10.1101/2021.05.28.446142

**Authors:** A.T. Hussin, S. Abbaspoor, K.L. Hoffman

**Affiliations:** Department of Biology, Centre for Vision Research, York University, Toronto, ON, Canada; Department of Psychology, Center for Integrative and Cognitive Neuroscience, Vanderbilt Vision Research Center, Vanderbilt Brain Institute, Vanderbilt University, Nashville, Tennessee

## Abstract

To understand the neural activity behind recollections of the distant past, we recorded from the hippocampus and retrosplenial cortex of macaques as they retrieved year-old and newly-acquired object-scene associations. Year-old memoranda preferentially evoked retrosplenial sigma oscillations (10-15 Hz), and phase-locking between RSC and eye movements during retrieval. In contrast, gamma-band retrosplenial-hippocampal synchrony was stronger during retrieval of new items. Primate retrosplenial oscillations may therefore guide retrieval of visuospatial events long past.

---

The ability to recall events that transpired months or years in the past is a crowning achievement of primate cognition, and one that is fundamental to our daily lives. Both the hippocampus and retrosplenial cortex are important structures for forming long-lasting memories of spatiotemporally distinct events (*1-2*). The neural mechanisms within these structures that give rise to retrieval over large time spans is unclear, though the retrosplenial cortex may assume a greater, more independent role over time (*3-9*). Tests of this role have been limited by i. species-adapted tasks in rodents that use shorter delays and restricted memorandum, ii. a historic emphasis on hippocampal physiology, and iii. indirect access to neuronal populations in humans. Here, we measure the neural activity in retrosplenial cortex and the hippocampus of macaques as they perform memory tasks on unique item-in-visuospatial-scenes. Some of these stimuli had not been seen for over a year, whereas other stimuli were presented for the first time and recalled within the same session or within 24 hours (Figure 1a). We asked whether the hippocampus (HPC) or retrosplenial cortex (RSC) would show changes in neural population activity as a function of retrieval of events learned 1-1.5 years earlier compared to newly-learned events.

**Fig. 1.**
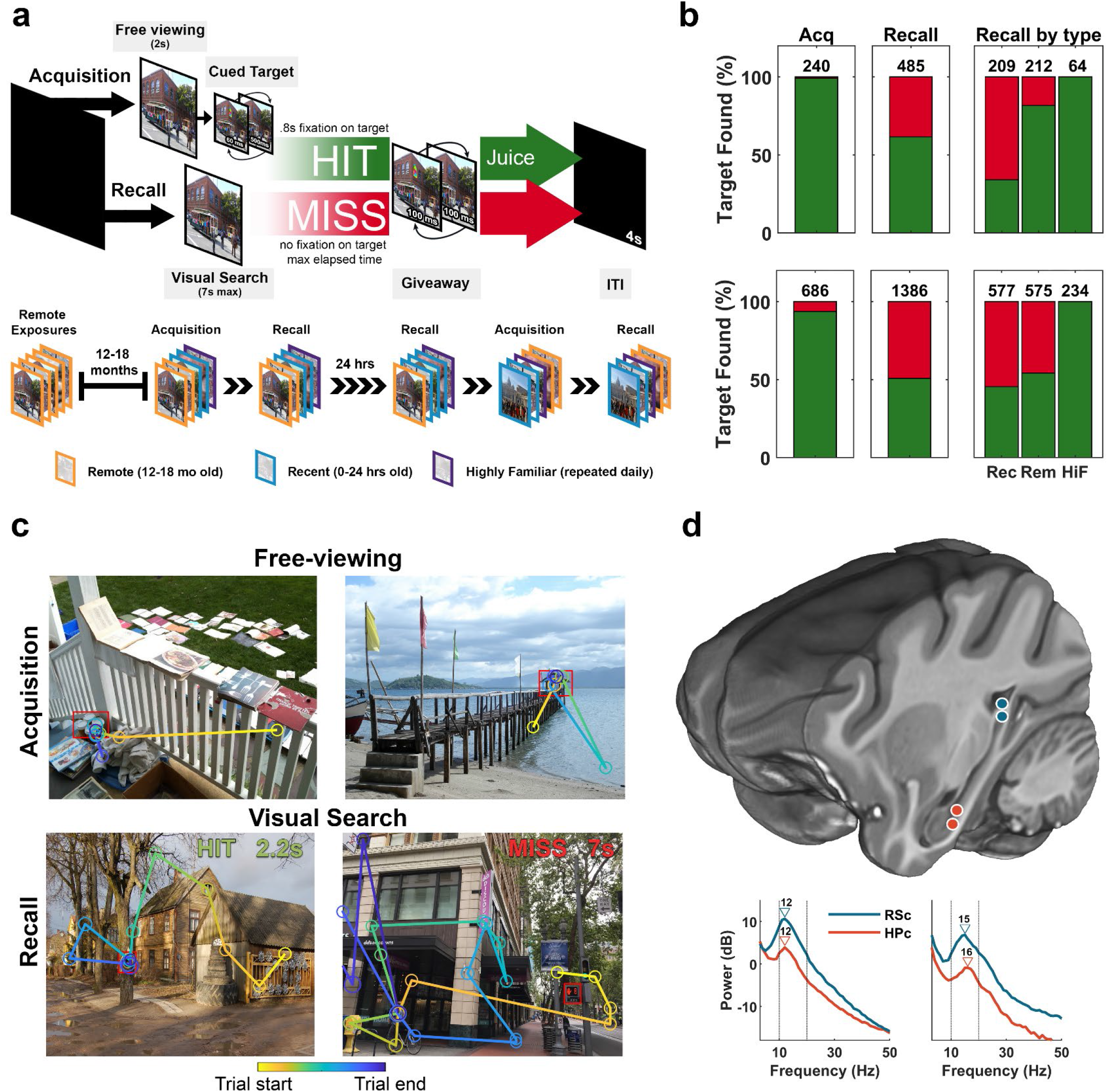
Experimental design and task performance. **a)** *Top*; acquisition trials begin with 2s of free viewing followed by target cueing. During recall, the scene is presented without this cue, requiring the target location to be remembered. The trial ends when gaze is maintained within the target area for 0.8s, for which a fluid reward is delivered (‘HIT’), or when the maximum trial time of 7s is reached (‘MISS’). A *giveaway* presents the cued target for a longer duration (100ms) at the end of a trial followed by an inter-trial interval of 4s. *Bottom*; sets were comprised of three stimulus types: ‘remote’ scenes, which had been presented 12-18 months prior, ‘recent’ scenes which were novel scenes at the start these sessions, and six ‘highly familiar’ scenes which were presented repeatedly throughout the study. Equal numbers of remote and recent scenes were presented in each set, randomly interleaved. Recall sets were shown after the acquisition set on the same day, and one day following acquisition, but before a new acquisition set. **b)** *Top*; from left, HIT rate for acquisition trials, recall trials, and recall by scene category, all for monkey RI. *Bottom*; same as top but for monkey LE. Values above bars indicate number of HITS in respective conditions. Rec = recent; Rem= remote; HiF = highly familiar. **c)** *Top row*: example scan paths during free viewing on remote acquisition trials. Outlined in red is the target. Directed gaze towards the (uncued) target indicates memory for the target. *Bottom row*: example scan paths for a HIT (remembered trial) and a MISS (forgotten trial) during recall trials. Top right on image shows search time in seconds. **d)** *Top:* Imaging-localized electrode positions, RSC in blue and HPC in red. *Bottom:* mean power during search for RSC (blue) and HPC (red; *left*: RI, *right*: LE). Vertical lines indicate 10 and 20 Hz; triangles indicate peak frequency.w

We recorded 62 sessions from two macaques (LE = 37, RI = 25), both of which had a >90% hit rate on acquisition trials. During recall, they had a higher hit rate for old (‘remote’) compared to newly acquired (‘recent’) scenes (Figure. 1b; LE: X^2^(1, 395) = 35, p<0.01; RI: X^2^(1, 395) = 44, p<0.01), indicating memory savings of remotely-learned scene targets. Using multichannel recordings from 12 indwelling, 16-channel linear electrodes, we obtained local field potentials from several structures, including the RSC and HPC (Figure 1c). The spectral power during visual search showed a prominent peak between 10-20 Hz in the RSC and HPC of both animals (Figure 1d). When we aligned the RSC signals to *Scene onset* and *Remembered- target selection* consisting of the last 1.5s before target selection on remembered trials, we found greater RSC power as early as 0.5s after scene onset (Figure 2b, c) and in the final second before target selection on remembered trials (Figure 2f, g), using multiple-comparison corrected permutation tests. We found no consistent differences in HPC power between remote and recent trials (Figure 2d and 2h), at any frequencies from 3-80 Hz. The anterior cingulate cortex (ACC) was recorded in one animal, showing oscillations in a higher, 23 Hz frequency band that was stronger for remote than recent trials, but only during scene onset (Extended Data Fig. 1).

**Fig. 2.**
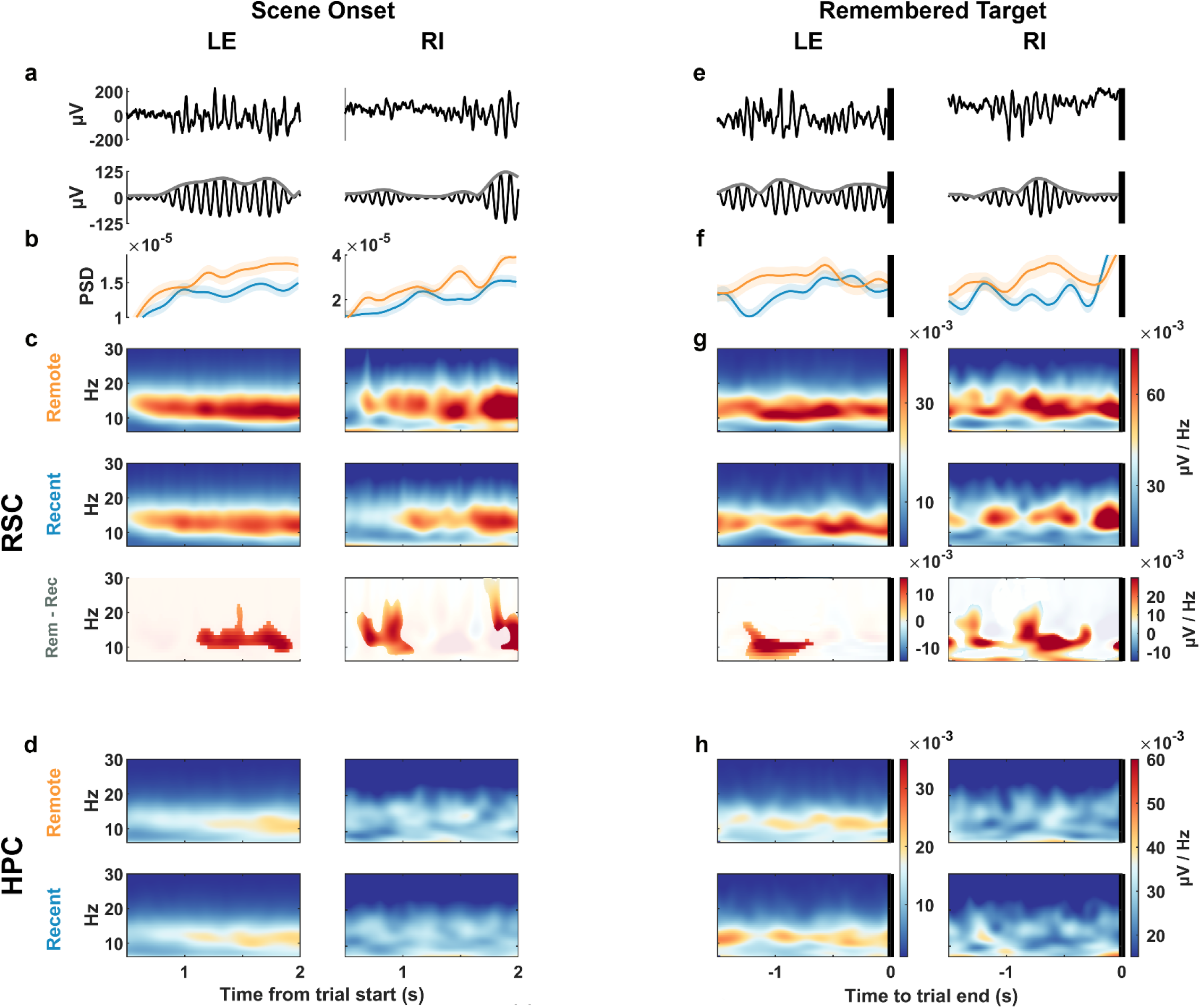
Differences in 10-15 Hz oscillations during recent and remote scene search. **a)** *Top*; Example broadband signal from RSC during the first 2s after scene onset. *Bottom*; 10-15 Hz filtered signal from above example. LE and RI indicate the respective study animals. Abscissas as in D. **b)** mean power spectral density of 10-15 Hz band using 500ms windows in 1ms steps. Shading indicates 95% bootstrap confidence intervals. **c)** *Top*; mean spectrogram of remote trials (LE, n = 594; RI, n = 152), middle; for recent trials (LE, n = 605; RI, n = 240), *bottom*; remote – recent difference spectrogram with the non-masked region representing areas with a difference of p < 0.05 in a cluster-based permutation test corrected for multiple comparisons. **d)** *Top*; mean spectrogram of remote trials in the hippocampus, *bottom*: recent. **e-h** same as **a-d** respectively but aligned to the last 1.5s before the end of remembered trials.

We used a general linear model (GLM) to quantify whether event age predicts magnitude of the sigma (10-15 Hz) RSC oscillation during acquisition trials, while accounting for other predictors, including the target location in visual search, and by animal. RSC sigma strength was predicted by memory age (remote>recent) and animal (RI>LE), but not target location (F (3,545) = 49.58, p = 2.2⨯10^−16^, adjusted-R^2^=0.21; age t=-3.08, p <0.01; animal t=11.92, p<0.001) Similarly, a GLM predicting RSC sigma on recall trials found that memory age (remote), and animal (RI) were significant predictors: (F (5,984) = 22.31, p=2.2⨯10^−16^, adjusted-R^2^=0.097; scene age (t=-4.07, p<0.001); animal (t=10.14, p<0.0001)) see supplementary methods for GLM details.

If memory-related retrosplenial cortex activity guides visual search during retrieval, one would expect greater coupling to the eye movements that underlie successful search, similar to previous observations with hippocampal signals (*10*). Specifically, we measured phase alignment of the LFP in both RSC and HPC to eye movements during successful retrieval, evaluated separately for remote and recent trials. Phase alignment was greater in the RSC (Figure 3b and 3e) on remote vs. recent scenes, beginning shortly before fixation onset (RI: -175 ms, LE: -75 ms) and lasting until 125-200 ms post-fixation. This effect was limited to remembered trials (Extended Data Fig. 2). In contrast to the RSC, the HPC phase alignment did not vary by scene age (Figure 3a and 3d). Thus, onto a general background of phase locking to visual fixations across regions, the retrosplenial cortex specifically showed greater coupling to the decision-making apparatus for old memory retrieval, consistent with a role for this region in memory-based guidance of actions.

**Fig. 3.**
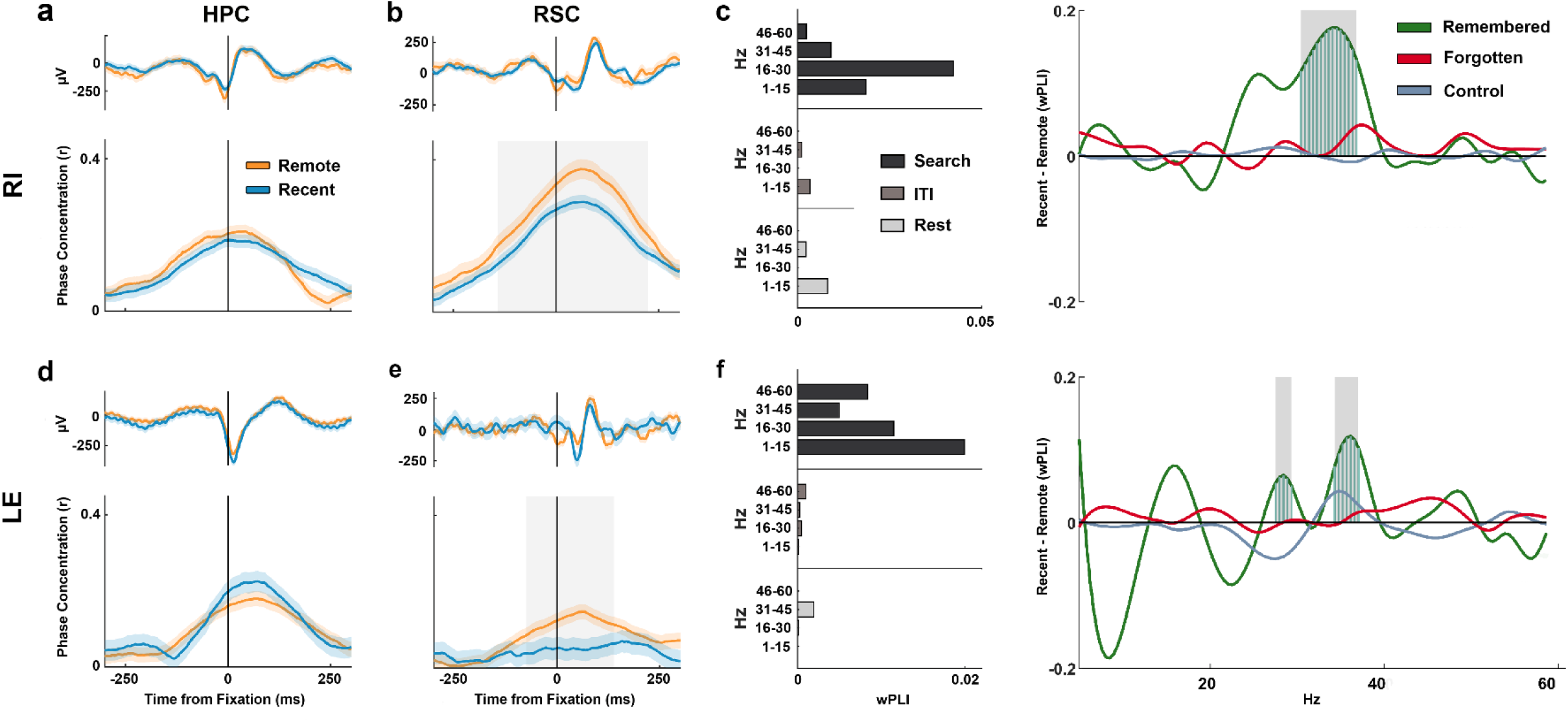
Retrosplenial cortex synchrony during retrieval includes phase-alignment with fixations during remote trials, and with HPC oscillations preferentially during recent trials. **a)** *Top*; mean LFP locked to fixations on remembered trials for both remote (orange) and recent (blue) scenes, *bottom*; mean phase concentration of HPC around the time of fixations for monkey RI (RI; remote n = 1297, recent n = 2972). Shading around mean traces represents 95% bootstrapped confidence intervals. **b)** Same as a) but for RSC (remote n = 1267, recent n = 2921). Grey shading represents p<0.05 difference between remote and recent phase concentrations in a two-tailed cluster-based permutation test. **c)** *Left*; phase-locking (wPLI) by frequency during 10-15 Hz RSC bouts across rest (n = 2338), ITI (n = 1937), search (n = 890) for RI. *Right*; difference in RSC-HPC synchrony during sigma bouts in recent and remote trials during remembered (recent = 106 bouts, remote = 176 bouts), forgotten (recent = 468, remote = 140) and low-sigma control bouts (recent = 1108, remote = 371). Vertical blue bars from zero and grey shading indicate frequencies where the difference between recent and remote is p<0.05 in a cluster-based permutation test. **d)** Same as a) but for monkey LE (remote n = 2054, recent n = 1031). **e)** same as b) but for monkey LE (remote n = 2134, recent n = 1042). **f)** Same as e) but for animal LE, *left*; rest (n = 3346), ITI (n = 6423) and search (n = 2384), *right*; remembered (recent = 112, remote = 204), forgotten (recent = 1159, remote = 909) and control (recent = 2937, remote = 2331).

In rats and mice, the neural ensembles in RSC coordinate their activity with hippocampal ensembles (*5, 9, 11, 12*), and these interactions may be essential for the laying down of long- lasting mnemonic representations. To assess interactions between these two structures, we calculated their debiased weighted phase lag index (wPLI) during sigma bouts obtained from search, inter-trial-intervals, and rest epochs. We found that the greatest phase synchrony in both animals occurred during search. We further examined inter-area synchrony during successful retrieval, comparing effects for remote and recent retrieval events. We found that during remembered, but not forgotten or low-sigma control trials, RSC-HPC synchrony was greater for recent compared to remote scenes, using a cluster-based permutation test corrected for multiple- comparisons (p<0.05). The only frequencies showing wPLI differences with memory age fell in the gamma range of ∼25-40 Hz (Figure 3c, f). This difference in synchrony was present only during recall trials and not during acquisition trials. Similarly, during the beta-bouts characteristic in the ACC, ACC-HPC synchrony for recent trials was stronger than for remote trials, but only in the ∼30 Hz range (Extended Data Fig. 3).

The prevalence and task-dependence of the 10-15 Hz sigma oscillation we observed in RSC was unexpected. The homologous structure in rodents, from which nearly all physiological evidence is based (but note (*13*)), is characterized by 7-8 Hz theta-band modulation (*3*), which appears to interact with the hippocampal theta oscillation (*11, 14*). In the relative absence of hippocampal theta oscillations during active search in the primate (*15*) it would appear, empirically, that the RSC exhibits a strong, band limited sigma oscillation. The closest match to this oscillatory phenotype appears during visual attention (*16*) apparently driven by pulvinar inputs (*17, 18, 19*). The primacy of vision in anthropoid species, and specifically the encoding of visuospatial metrics within structures supporting spatial exploration and memory (*2*) may result in a greater influence of visual cortex and thalamus on these structures. This may, in turn, be integrated with a phylogenetically conserved role for the RSC in gradual development of representations over time (*8, 20*). The connectivity of RSC in macaques situates it as proximate to other regions critical for visuospatial processing and for spatiotemporal event memory (*3*). In light of the paucity of neural data on old memories in primates, and the prominence of the neural oscillations we observed, the present findings offer a strong guidance for future neural data analysis of mnemonic content within the RSC and information routing to other structures, as memories age.

## Methods

### Surgical procedures

Two adult female macaques (*Macaca mulatta*, “LE” and “RI”, approximately 12 and 10 kg, respectively) were implanted in a sterile surgical procedure with 12 indwelling flexible, polyamide-based intracortical 16-channel linear electrode arrays with contacts at 130 µm spacing (prototypes to the ‘Microflex’, Blackrock electrodes), targeting the hippocampus, anterior cingulate and retrosplenial cortices as (described in (*21*)). All surgical and experimental protocols were conducted with approval from the local ethics and animal care authorities (Animal Care Committee, Canadian Council on Animal Care). Surgery was performed and data were collected at York University, Toronto, Canada.

### Task design

Both monkeys completed a memory-guided visual search task 12-18 months prior to the present recordings. During this task, a target object was embedded in a naturalistic scene, and presented alongside other objects-in-scenes, comprising the ‘remote’ stimuli used in this study. During the present experiments, the animals performed two task versions within each daily session: ACQUISITION and RECALL trials. During an acquisition trial, the scene is displayed for 2s and the animal is allowed to view the scene freely, followed by presentation of a target unique to the scene that is cued by alternating between original (500ms) and complementary colors (60ms), making the target location salient and appearing to “pop out” to the observer. Target cueing began at 2s and continued until the target was selected (designated as a HIT) or until 7s had passed (designated as a MISS). Selection of the target was accomplished by holding gaze in the target region for a prolonged duration (≥800ms). Once the target was cued, the task became trivially easy for the macaques. In contrast, during a recall trial, the scene is presented without the cue and the animal had 7s to find and select the (un-cued) target for juice reward (HIT or remembered) or the trial ended without reward (MISS or forgotten). All trials ended with a giveaway, where the original and colour-modified scene alternate (100ms each x 5) revealing the target to the animal. An inter-trial interval of 4s of black screen followed each trial (Figure 1A).

Scenes were grouped into sets of 12 (monkey RI) or 16 (monkey LE) scenes. The number of scenes in a set was designed to account for individual performance differences. Each set had three types of scenes; *recent, remote* and *highly familiar*. Recent scenes were novel to the animal during the first acquisition presentation. Remote scenes were scenes used during initial task training 12-18 months prior. Highly familiar scenes were a preselected subset of six remote scenes that were repeated regularly throughout the experiment, and therefore have a high HIT rate. Two of these six scenes were included in each set. Presentation order of scene types was randomized within the set. Recall sets followed immediately the acquisition sets within a daily session. In addition, the following day’s session began with the recall sets from the previous day. Two new sets were presented each day (i.e. 24 and 32 new scenes per day, for the two animals, respectively). Daily sessions started and ended with a 5-minute rest period where a black screen was presented. Eye movements were recorded at 1250 Hz using video-based eye tracking (iViewX Hi-Speed Primate Remote Infrared Eye Tracker). For the analysis we excluded trials where the animals spent >20% of trial duration looking off-screen (monkey LE: 238/2755 or 8%, monkey RI: 50/1310 or 4%), to ensure that only trials where the animals were attending to the task were included.

### Neural recordings

Local-field potentials (LFP) were recorded simultaneously from the hippocampus, anterior cingulate and retrosplenial cortices, digitally sampled at 32 kHz using a Digital Lynx acquisition system (Neuralynx, Inc.) and filtered between 0.5 Hz and 2 kHz. Due to problems with the implant connection to the anterior cingulate cortex (ACC) contacts of one animal, results from ACC were evaluated for only the remaining animal (RI). No well-isolated units were recorded with these arrays, so all analysis is based on local field potentials. Inclusion criteria for any probe consisted of CT/MR coregistration or post-explant MR verification of location within an ROI, including visualization of marker lesions, for the subset of probes in which they had been delivered. Along the 16-channel probe, occasional channels were excluded based on signal not following biological 1/f spectra or otherwise not showing signal. The remaining neural signals were downsampled to 1 kHz, and a notch filter (59.9 to 60.1 Hz) was used to remove 60 Hz noise. All offline behavioral and neural analysis was conducted in MATLAB using custom-written scripts and FieldTrip ((*22*); fieldtrip.fcdonders.nl).

### Generalized eigendecomposition (GED)

We implemented generalized eigendecomposition (GED) for source separation of the LFP in the multichannel arrays (based on methodology described in *23*)), due to probes’ non-orthogonal alignment to laminae, and our inclusion of multiple probes within an area. For analyses of power, phase concentration and phase synchrony, this involved the use of linear spatial filters to provide a weighted combination of electrode activity, isolating sources of independent variance in multichannel data. The spatial filters were defined by the generalized eigendecomposition (GED) of channels’ covariance matrices. The two separate covariance matrices selected will result in eigenvectors that maximally differentiate them. If the signal features to be accentuated and those to be attenuated are designated by S and R respectively, the eigendecomposition problem can be written as SW = WRΛ. The solution of this problem yields W which is a matrix of eigenvectors and Λ that is a diagonal matrix of eigenvalues. The resultant filters, defined by eigenvectors, are then applied to the multichannel electrode time series to obtain a set of component time series. If GED was unable to differentiate between various sources of variance, shrinkage regularization was employed at 1 percent.

For power and phase concentration analyses, the S matrix was created from 1 second of signal after scene onset (start of trial) and the R matrix from 1 second of baseline activity prior to the scene onset. In this design, we sought to attenuate continuous noise in the signal, and accentuate the dynamics and signal sources producing those dynamics that are relevant to the task. For phase synchrony analysis, we created the S matrix from the band-pass filtered electrode time series in 10-20Hz. The R matrix was then formed from the broadband electrode time series. In this case, the column in W with the highest corresponding eigenvalue then corresponds to the eigenvector that maximally enhances the 10-20 Hz frequency activity. The inputs to the GED were signal from multiple channels and trials from a given probe and the output was a single weighted time-series component per trial per probe. The analyses that follow use the primary resultant GED component that represents the weighted combination of activity from multiple channels in each probe. This source isolation method circumvents the tradeoff of excluding simultaneously-recorded channels arbitrarily, when they may contribute informative signal, while ensuring that the dependence of signals across channels is not falsely considered as independent samples of the timeseries. Qualitatively, use of single-channel LFP yielded very similar results, though it suffers from the aforementioned arbitrary exclusion issue.

### Spectral analysis

Grand power was computed using a Fourier transform and a Hanning multi-taper frequency transformation, averaging over the whole duration of search trials including both acquisition and recall trials (N trials for LE = 1152, RI = 422). Mean power spectral density was examined in 500ms windows with a 1ms sliding window conducted on individual trials then averaged across trials. For mean time-frequency spectra, we implemented a Morlet wavelets multi-taper transformation with a width of five cycles and a frequency step-size of 1 Hz.

### Phase concentration

To examine phase alignment with eye movement we inspected the LFP signal in 600ms windows centered around fixation onsets (peri-fixation signal). We examined all recall trial fixations split by remembered and forgotten trials. We bandpass filtered the peri-fixation neural signal between 4-9 Hz, then used the phase angles of the Hilbert transform to compute the mean resultant vector length (or phase concentration). Circular statistical analyses were performed using the Circular Statistics Toolbox for MATLAB (*24*).

### Bout detection

For detection of oscillatory bouts of activity in the dominant frequency band, the RSC signal from all trials was bandpass filtered between 10-15 Hz. Bouts were defined as time periods in which the signal exceeded a threshold of 2 SDs above the envelope mean, and for a minimum duration of 100ms above 1 SD. Bout amplitude was defined as the maximum of the envelope within a bout. Control bouts were chosen under the same criteria but in opposite direction (i.e. -2 SD) to identify windows of time with the weakest RSC 10-15 Hz power.

### Phase synchrony

Inter-areal phase synchrony during bouts was calculated from the cross-spectral density of the RSC signal and the corresponding HPC signal using the debiased weighted phase lag index (wPLI). The debiased wPLI measure of phase-synchronization minimizes the influence of volume-conduction, noise and the sample-size bias (*25*), allowing us a more conservative measure of synchrony in that it reduces our Type 1 error, while recognizing that zero-lag effects may go undetected (Type II error).

### Statistical analysis

Proportions of hit rate and bout occurrence across scene types were compared using a two-tailed Chi-square test for comparing proportions. Search times were compared using a two-tailed rank-sum Wilcoxon test. Time-frequency spectra were compared using nonparametric permutation tests using the Monte Carlo sampling method and a cluster-based correction for multiple-comparisons. To test for statistical significance of differences between phase concentration and synchrony (wPLI values) during the recent and remote conditions, we performed a nonparametric permutation test with the difference in phase concentration or coherence between conditions as our test statistic. The test statistic was calculated for each frequency bin, then bins whose statistic value was <2.5th or >97.5th percentiles were selected, and cluster-level statistics were calculated by summing the test statistic within a cluster. This testing method corresponds to a two-tailed test with false-positive rate of 5% corrected for multiple comparisons across frequencies (*26-28*).

## Acknowledgments

The authors would like to acknowledge the animal care and training support from Patricia Sayegh and Natasha Down, and image registration pipeline development from Wolf Zinke.

## Author contributions

A.T.H. and K.L.H. conceived, designed, interpreted the study, and wrote the manuscript; A.T.H. collected the data; A.T.H., S.A. and K.L.H. analyzed the data.

## Data and materials availability

All data, code, and materials used in the analysis will be made available upon request for purposes of reproducing or extending the analysis.

**Extended Data Fig. 1.**
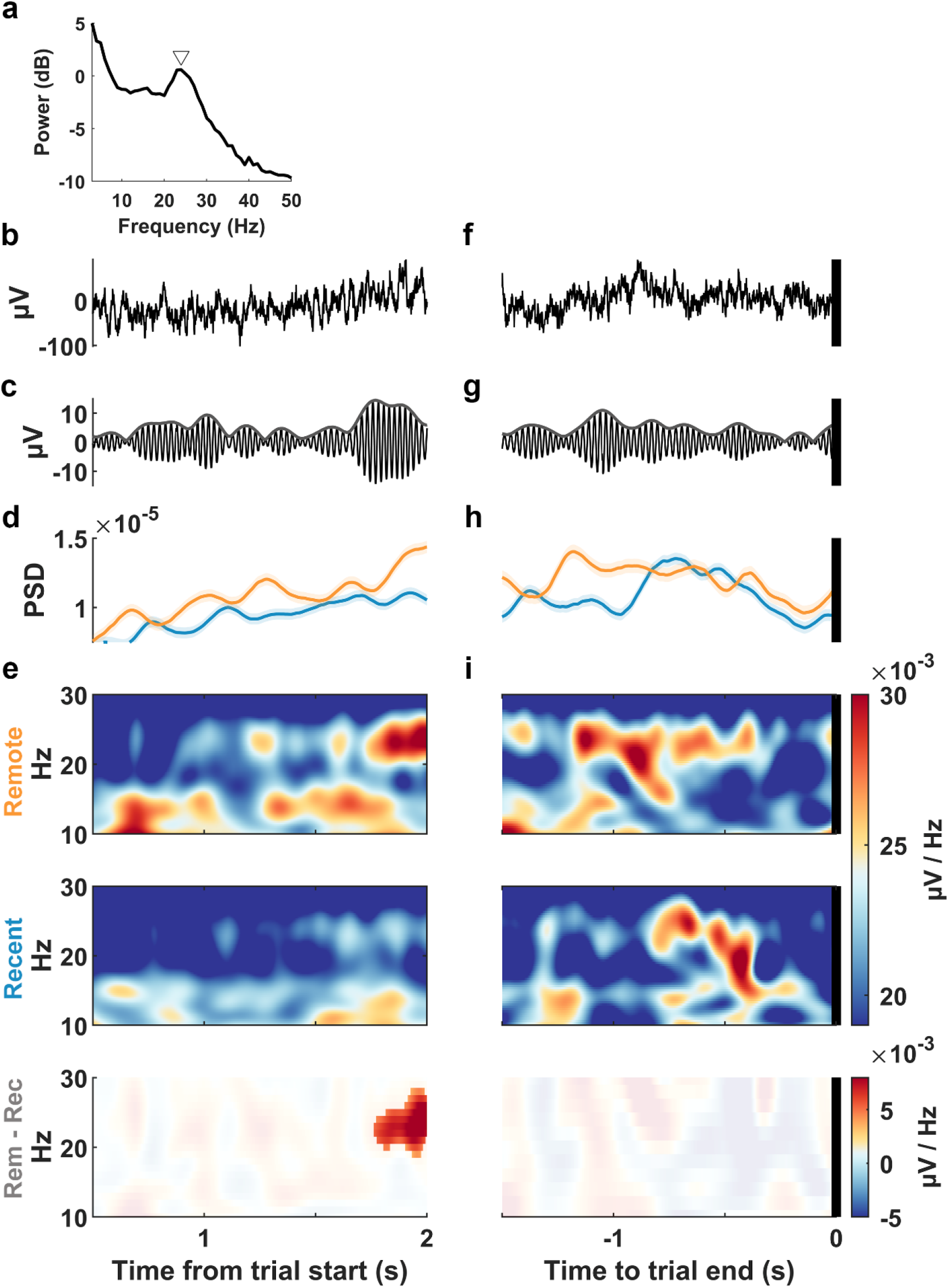
Anterior cingulate cortex (ACC) exhibits greater 21-26 Hz power during remote scenes. a) Power spectrum during search showing a peak around ∼23 Hz indicated by triangle. b) Example broadband signal during first 2s after trial onset when a scene is presented. c) 21-26 Hz filter of signal above. d) mean power spectral density of 21-26 Hz band using 300ms windows in 1ms steps. Shading indicates 95% bootstrap confidence interval. e) Top; mean spectrogram of remote trials (n = 289), middle; recent trials (n = 193), bottom; remote – recent difference spectrogram with the non-greyed region representing cluster with a difference of p < 0.05 in a permutation test. f-i) same as b-e but for trial end epoch of remembered trials (remote n = 45, recent n = 65).

**Extended Data Fig. 2.**
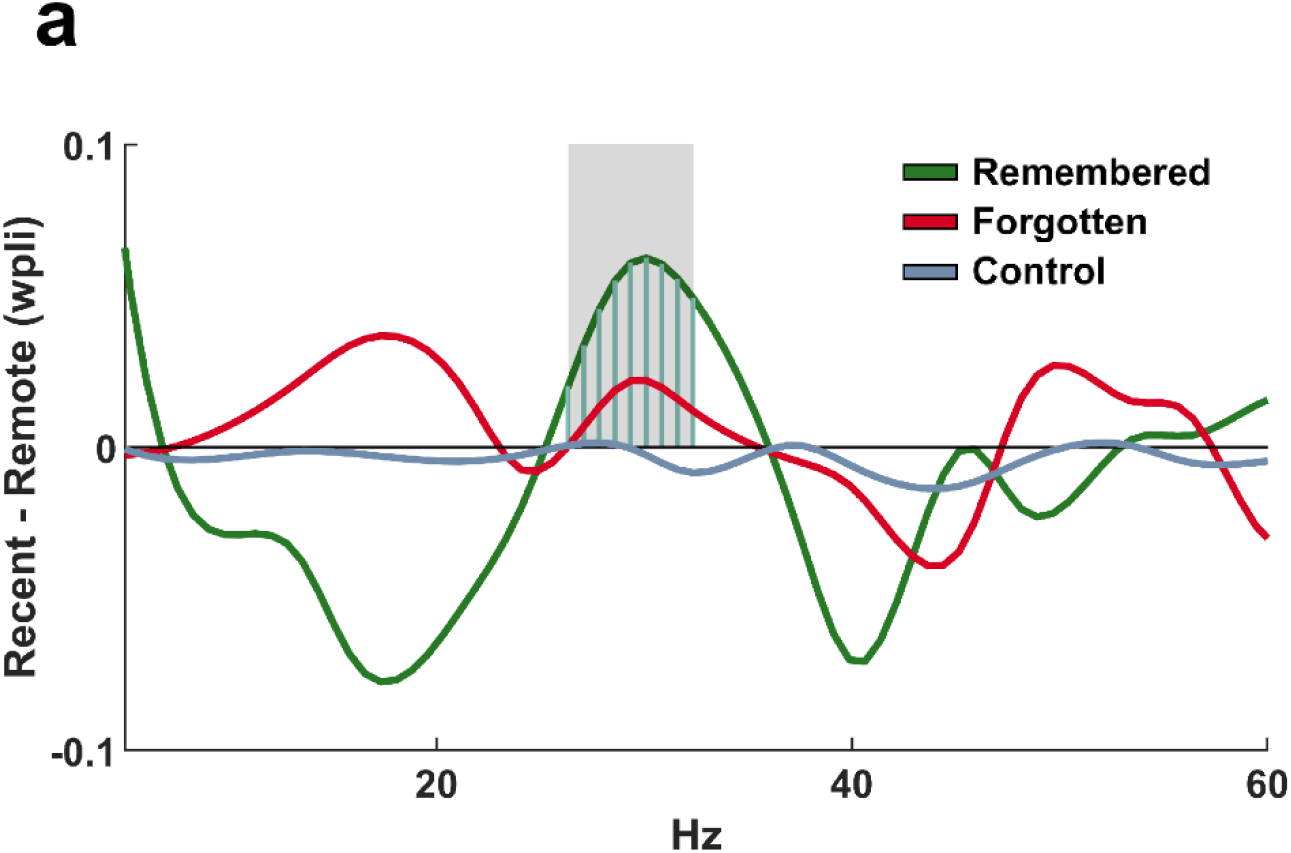
Greater phase synchrony between anterior cingulate cortex (ACC) and hippocampus (HPC) during remembered recent than remote scenes. a) difference in ACC-HPC phase locking between bouts on recent and remote trials across remembered (recent = 111 bouts, remote = 194 bouts), forgotten trials (recent = 530, remote = 142) and control bouts (recent = 405, remote = 718). Vertical blue bars from zero indicate frequencies with a permutation difference of p<0.05.

**Extended Data Fig. 3.**
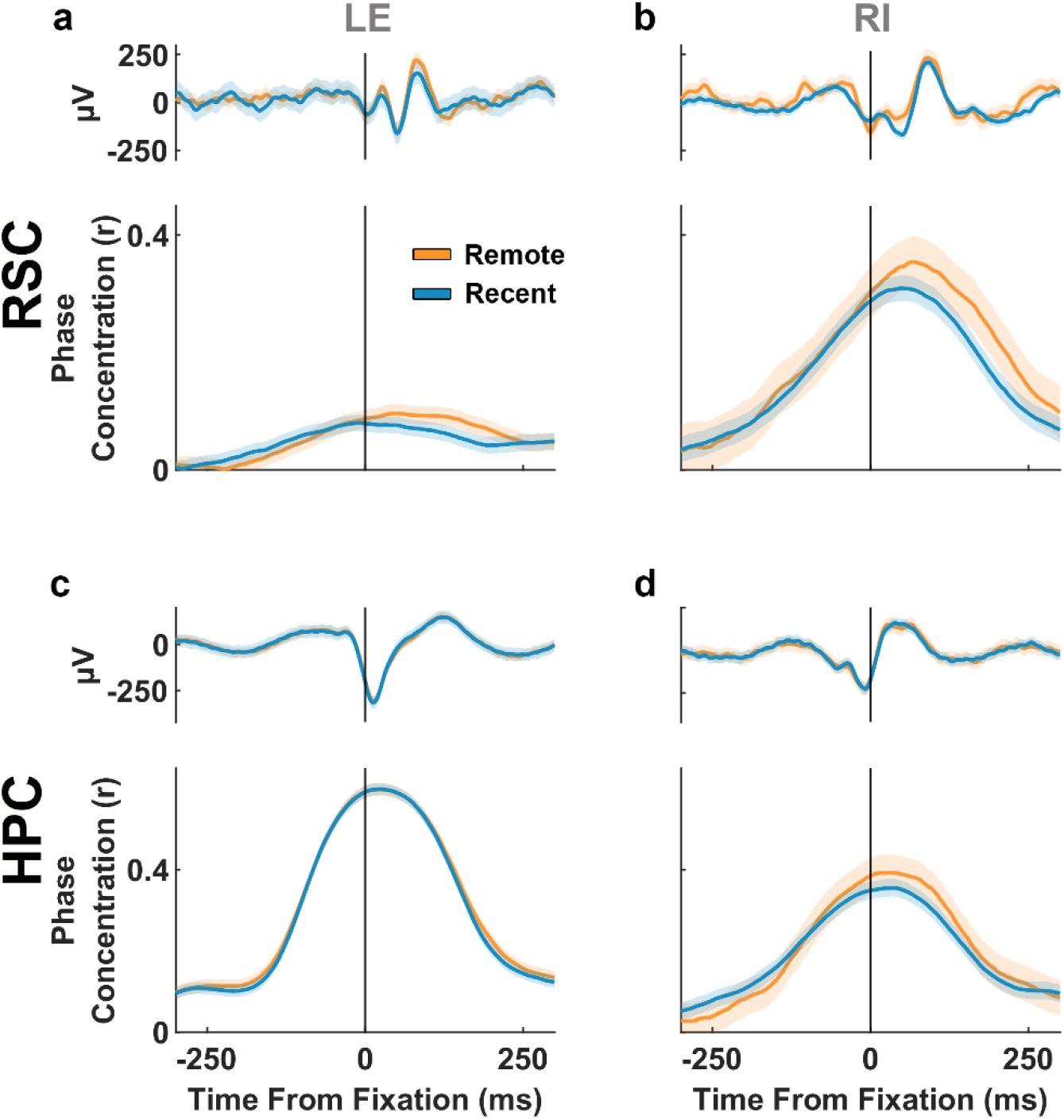
During forgotten trials, neither retrosplenial cortex nor the hippocampus show differences in peri-fixation phase locking for remote compared to recent trials. a) Top; mean LFP locked to fixations on forgotten trials, bottom; mean phase concentration of retrosplenial cortex (RSC) around fixations for monkey LE (remote n = 2134, recent n = 1042). Light shade around mean traces represent 95% bootstrapped confidence intervals. b) same as a) but for animal RI (remote n = 1267, recent n = 2921). c) and d) are similar to a) and b) but for the hippocampus (HPC) of each animal respectively.

